# Transposon silencing in the *Drosophila* female germline ensures genome stability in progeny embryos

**DOI:** 10.1101/363028

**Authors:** Zeljko Durdevic, Ramesh S. Pillai, Anne Ephrussi

**Affiliations:** Developmental Biology Unit, European Molecular Biology Laboratory, Meyerhofstrasse 1, Heidelberg, D-69117, Germany; Department of Molecular Biology, University of Geneva, 30, quai Ernest-Ansermet, Geneva 4, CH-1211, Switzerland

**Keywords:** Vasa, transposons, DNA damage, Chk2, embryogenesis

## Abstract

The piRNA pathway functions in transposon control in the germ line of metazoans. The conserved RNA helicase Vasa is an essential piRNA pathway component, but has additional important developmental functions. Here we address the importance of Vasa-dependent transposon control in the *Drosophila* female germline and early embryos. We find that transient loss of *vasa* expression during early oogenesis leads to transposon up-regulation in supporting nurse cells of the fly egg-chamber. We show that elevated transposon levels have dramatic consequences, as de-repressed transposons accumulate in the oocyte where they cause DNA damage. We find that suppression of Chk2-mediated DNA damage signaling in *vasa* mutant females restores oogenesis and egg production. Damaged DNA and up-regulated transposons are transmitted from the mother to the embryos, which sustain severe nuclear defects and arrest development. Our findings reveal that the Vasa-dependent protection against selfish genetic elements in the nuage of nurse cell is essential to prevent DNA damage-induced arrest of embryonic development.

## Introduction

Transposons and other selfish genetic elements are found in all eukaryotes and comprise a large fraction of their genomes. Although transposons are thought to be beneficial in driving evolution (Levin and Moran, 2011), their mobilization in the germline can compromise genome integrity with deleterious consequences: insertional mutagenesis reduces the fitness of the progeny and loss of germ cell integrity causes sterility. Therefore, it is of great importance for sexually reproducing organisms to firmly control transposon activity in their germ cells. Metazoans have evolved a germline specific mechanism that, by relying on the activity of PIWI family proteins and their associated Piwi-interacting RNAs (piRNAs), suppresses mobile elements.

*Drosophila* harbors three PIWI proteins: Piwi, Aubergine (Aub) and Argonaute 3 (Ago3) which, guided by piRNAs, silence transposons at the transcriptional and posttranscriptional levels (reviewed in Guzzardo et al., 2013). Besides PIWI proteins, other factors such as Tudor-domain proteins and RNA helicases are involved in piRNA biogenesis and transposon silencing. Mutations in the majority of piRNA pathway genes in *Drosophila* females cause transposon up-regulation that leads to an arrest of oogenesis. This effect can be rescued by suppression of the DNA damage checkpoint proteins of the ATR/Chk2 pathway (Chen et al., 2007; Klattenhoff et al., 2007; Pane et al., 2007). In contrast, inhibition of DNA damage signaling cannot restore embryonic development (Chen et al., 2007; Klattenhoff et al., 2007; Pane et al., 2007). Recent studies suggest that PIWI proteins might have additional roles during early embryogenesis independent of DNA damage signaling (Khurana et al., 2010; Mani et al., 2014). However, functions of the piRNA pathway during early embryonic development remain poorly understood.

One of the essential piRNA pathway factors with an important role in development is the highly conserved RNA helicase Vasa. First identified in *Drosophila* as a maternal-effect gene (Hay et al., 1988; Lasko and Ashburner, 1990; Schüpbach and Wieschaus, 1986), *vasa* (*vas)* was subsequently shown to function in various cellular and developmental processes (reviewed in Lasko, 2013). In the *Drosophila* female germline Vasa accumulates in two different cytoplasmic electron-dense structures: the pole (or germ) plasm at the posterior pole of the oocyte, and the nuage, the perinuclear region of nurse cells. In the pole plasm, Vasa interacts with the pole plasm inducer Oskar (Osk) (Breitwieser et al., 1996; Jeske et al., 2015) and ensures accumulation of different proteins and mRNAs that determine primordial germ cell (PGC) formation and embryonic patterning (Hay et al., 1988; Lasko and Ashburner, 1990). In the nuage, Vasa is required for the assembly of the nuage itself (Liang et al., 1994; Malone et al., 2009) and facilitates the transfer of transposon RNA intermediates from Aub to Ago3, driving the piRNA amplification cycle and piRNA-mediated transposon silencing (Nishida et al., 2015; Xiol et al., 2014). As Vasa’s involvement in many cellular processes renders it difficult to analyze its functions in each process individually, it remains unknown whether Vasa’s functions in development and in the piRNA pathway are linked or independent.

In this study, we address the role of Vasa in transposon control in *Drosophila* development. We find that failure to suppress transposons in the nuage of nurse cells causes DNA double-strand breaks (DSBs), severe nuclear defects, and lethality of progeny embryos. Even transient interruption of Vasa expression in early oogenesis de-represses transposons and impairs embryo viability. Depletion of the *Drosophila Chk2* ortholog *maternal nuclear kinase* (*mnk*) restores oogenesis in *vas* mutants, but does not suppress defects in transposon silencing or DSB-induced nuclear damage and embryonic lethality. We show that up-regulated transposons invade the maternal genome, inducing DNA DSBs that, together with transposon RNAs and proteins, are maternally transmitted and consequently cause embryogenesis arrest. Our study thus demonstrates that Vasa function in the nuage of *Drosophila* nurse cells is essential to maintain genome integrity in both the oocyte and progeny embryos, ensuring normal embryonic development.

## Results

### Vasa dependent transposon control is not essential for oogenesis

Vasa is required for piRNA biogenesis and transposon silencing in *Drosophila*, as in *vas* mutants piRNAs are absent and transposons are up-regulated (Czech et al., 2013; Handler et al., 2013; Malone et al., 2009; Vagin et al., 2004; Zhang et al., 2012). To investigate the importance of transposon control in *Drosophila* development, we expressed wild-type GFP-Vasa fusion protein (GFP-Vas^WT^; Supplementary Figure S1A) in the female germline of loss-of-function (*vas^D1/D1^*) *vas* flies using two promoters with distinct expression patterns (Supplementary Figure S1B-C): the *vas* promoter is active at all stages of oogenesis, whereas the *nos* promoter is inactive between stages 2 and 6 (Supplementary Figure S1B-C).

We first assessed the ability of GFP-Vas^WT^ fusion protein to promote transposon silencing in the female germline, and examined the effect of GFP-Vas^WT^ on the level of expression of several transposons in *vas* mutant ovaries. We chose the long terminal repeat (LTR) retrotransposons *burdock* and *blood*, and the non-LTR retrotransposon *HeT-A*, which were previously reported to be up-regulated upon Vasa depletion (Czech et al., 2013; Vagin et al., 2004). The LTR retrotransposon *gypsy*, which belongs to the so-called somatic group of transposons and is not affected by *vasa* depletion, served as a negative control (Czech et al., 2013). Loss-of-function *vas^D1/D1^* ovaries contained elevated levels of *burdock*, *blood* and *HeT-A* RNA (Figure 1A). Remarkably, silencing of transposons by GFP-Vas^WT^ in *vas^D1/D1^* flies depended on which Gal4 driver was used (Supplementary Figure S1B-C): when driven by *nos-Gal4*, GFP-Vas^WT^ had no effect on transposon levels, whereas when driven by *vas-Gal4* it led to the re-silencing of transposons (Figure 1A). This differential effect presumably reflects the stages of oogenesis at which the *nos* and *vas* promoters are active, and suggests that lack of Vasa between stages 2 and 6 of oogenesis (Supplementary Figure S1B) leads to transposon de-repression. Importantly, independent of Gal4 driver used, expression of GFP-Vas^WT^ restored oogenesis (Figure 1B and Supplementary Figure S1D) and egg-laying (Supplementary Figure S1E-F). The fact that in spite of transposon up-regulation oogenesis and egg-laying rates were fully restored in *vas^D1/D1^* flies (Figure 1A, indicated by + and – and Supplementary Figure S1D-F) indicates that oogenetic processes are not affected below a certain threshold of transposon activity.

**Figure 1.**
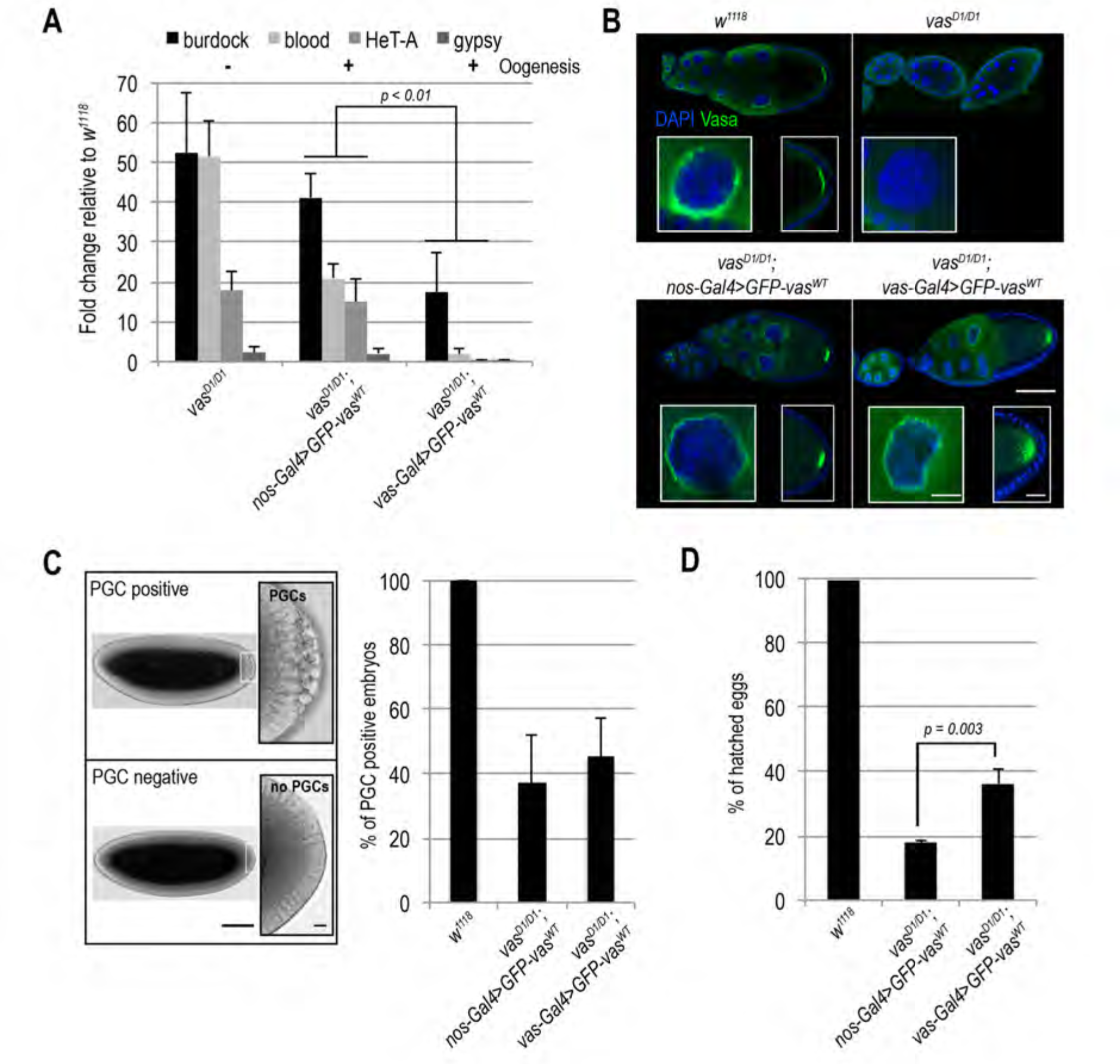
Silencing of transposon RNAs during oogenesis is essential for embryonic development. A) Q-PCR analysis of LTR transposons *burdock*, *blood* and *gypsy* and non-LTR transposon *HeT-A* RNAs in *vas^D1/D1^*, *vas^D1/D1^*; *nos-Gal4*>GFP-Vas^WT^ *and vas^D1/D1^*; *vas-Gal4*>GFP-Vas^WT^, ovaries. Expression level of transposons in wild-type (*w^1118^)* was set to 1 and normalized to *rp49* mRNA in individual experiments. Error bars represent the standard deviation among three biological replicates. *P*-values were determined by Student’s *t-*test. *P*-values for *burdock* (0.006), *blood* (0.0002) and *HeT-A* (0.0007) were lower than 0.01 (indicated in the chart), while *gypsy* levels were not significantly different (*p* = 0.5). Oogenesis completion is indicated with + and –. B) Immunohistochemical detection of Vasa in wild-type (*w^1118^)* and *vas^D1/D1^* flies (upper panel), and GFP signal of GFP-Vas^WT^ fusion protein in *vas^D1/D1^*; *nos-Gal4*>GFP-Vas^WT^ *and va^D1D/D1^*; *vas-Gal4*>GFP-Vas^WT^ flies (lower panel). Insets show enlarged images of nuage and oocyte posterior pole. Scale bars indicate 50 µm (egg-chambers) and 10 µm (nuage and pole plasm). C) Quantification of PGC positive embryos produced by wild-type (*w^1118^)*, *vas^D1/D1^*; *nosGal4*>GFP-Vas^WT^ *and vas^D1/D1^*; *vas-Gal4*>GFP-Vas^WT^ flies. Error bars represent the standard deviation among three biological replicates (Supplementary Table S4). Panel (left) shows PGC positive embryo (top) and PGC negative embryo (bottom). Scale bars indicate 100 µm (embryo) and 5 µm (PGCs). D) Hatching rates of eggs laid by wild-type (*w^1118^*), *vas^D1/D1^*; *nos-Gal4*>GFP-Vas^WT^ *and vas^D1/D1^*; *vas-Gal4*>GFP-Vas^WT^ flies. Error bars represent the standard deviation among three biological replicates (Supplementary Table S2). *P*-value was determined by Student’s *t-*test.

### Loss of Vasa during early oogenesis affects viability of progeny embryos

Concentration of Vasa protein at the posterior pole of the embryo is essential for PGC and abdomen formation during embryogenesis (Hay et al., 1988; Lasko and Ashburner, 1990; Schüpbach and Wieschaus, 1986). We analyzed number of PGC positive embryos and the hatching rate of eggs produced by *vas^D1/D1^* flies expressing GFP-Vas^WT^ either under control of the *nos* or the *vas* promoter (*vas^D1/D1^*; *nos-Gal4*>GFP-Vas^WT^ and *vas^D1/D1^*; *vas-Gal4*>GFP-Vas^WT^ embryos). PGC formation was restored in approximately 50% of *vas^D1/D1^*; *nos-Gal4*>GFP-Vas^WT^ and *vas^D1/D1^*; *vas-Gal4*>GFP-Vas^WT^ embryos (Figure 1C). However, DAPI staining revealed nuclear damage in some *vas^D1/D1^*; *nos-Gal4*>GFP-Vas^WT^ embryos (see below), which we excluded from the quantification.

Expression of GFP-Vas^WT^ also partially rescued the hatching of eggs produced by *vas^D1/D1^* flies (Figure 1D). However, expression of GFP-Vas^WT^ led to a significantly lower hatching rate in *vas^D1/D1^*; *nos-Gal4*>GFP-Vas^WT^ than in *vas^D1/D1^*; *vas-Gal4*>GFP-Vas^WT^ flies (Figure 1D). Expression of GFP-Vas^WT^ in heterozygous loss-of-function *vas^D1/Q7^* females led to a low hatching rate similar to *vas^D1/D1^* (Supplementary Figure S2A-B), excluding a possible secondary mutation as the cause of the low hatching rate. The fact that in spite of comparable GFP-Vas^WT^ levels (Supplementary Figure S2C) the hatching rate of *vas^D1/D1^*; *vas-Gal4*>GFP-Vas^WT^ embryos was higher than that of *vas^D1/D1^*; *nos-Gal4*>GFP-Vas^WT^ embryos, suggests that transient loss of *vas* expression during early oogenesis impairs viability of progeny embryos (Figure 1D).

### Elevated transposon levels cause DNA and nuclear damage in progeny embryos

Elevated transposon activity leads to DNA damage and ultimately to cell death. During our analysis of PGC formation we observed nuclear damage in a considerable fraction of *vas^D1/D1^*; *nos-Gal4*>GFP-Vas^WT^ embryos. Quantification revealed a high proportion of embryos with nuclear defects among *vas^D1/D1^*; *nos-Gal4*>GFP-Vas^WT^ embryos (Figure 2A). Transposon mobilization causes DSBs in genomic DNA that are marked by the incorporation of a phosphorylated form of the H2A variant (γH2Av), a histone H2A variant involved in DNA DSB repair. Analysis of γH2Av occurrence showed that embryos displaying nuclear damage were γH2Av-positive (Figure 2B), indicating that DNA DSBs cause nuclear defects. The levels of γH2Av were higher in *vas^D1/D1^*; *nos-Gal4*>GFP-Vas^WT^ embryos compared to wild-type and *vas^D1/D1^*; *vas-Gal4*>GFP-Vas^WT^ (Figure 2C).

**Figure 2:**
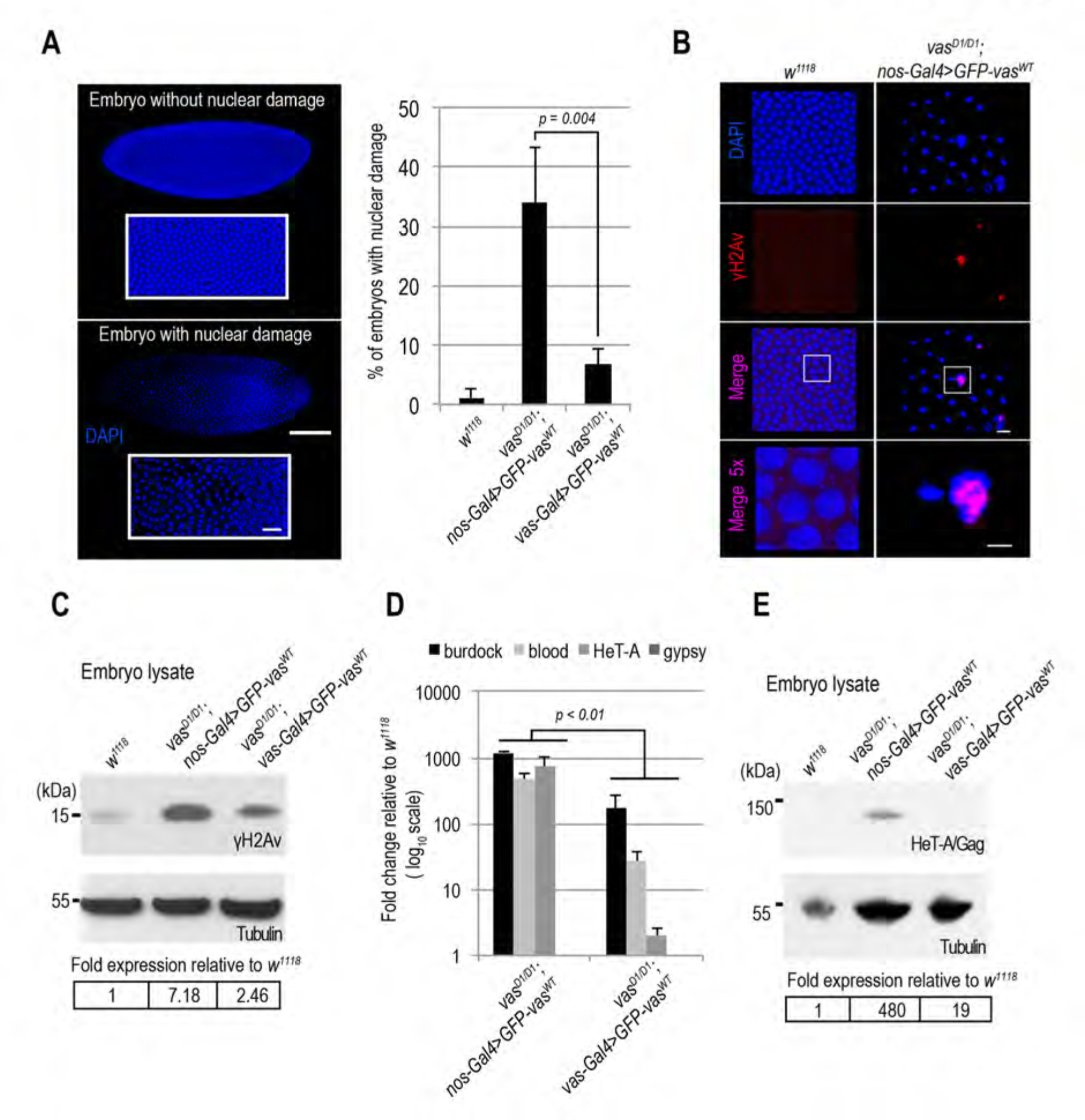
Maternally transmitted transposon RNAs cause DNA double strand breaks and nuclear damage in progeny embryos. A) Quantification of nuclear damage determined by NucBlue Fixed Cell Stain staining of wild-type (*w^1118^*), *vas^D1/D1^*; *nos-Gal4*>GFP-Vas^WT^ *and vas^D1/D1^*; *vas-Gal4*>GFP-Vas^WT^ stage 5 embryos. Error bars represent the standard deviation among three biological replicates (Supplementary Table S5). *P*-value was determined by Student’s *t-*test. Panel shows an embryo without (top) and an embryo with nuclear damage (bottom). Scale bars indicate 100 µm (embryo) and 10 µm (magnification). B) Immunohistochemical detection of DNA double strand breaks using antibodies against H2Av pS137 (γH2Av) in wild-type (*w^1118^*,) and *vas^D1/D1^*; *nos-Gal4*>GFP-Vas^WT^ stage 5 embryos. Whole embryos are presented in A. Scale bars indicate 5 µm and 2 µm (5x magnification). C) Western blot analysis using antibodies against H2Av pS137 (γH2Av) showing protein levels in wild-type (*w^1118^*), *vas^D1/D1^*; *nos-Gal4*>GFP-Vas^WT^ *and vas^D1/D1^*; *vas-Gal4*>GFP-Vas^WT^ 1-3 hour old embryos. Tubulin was used as a loading control. Table shows quantification of γH2Av levels relative to wild-type. gH2Av signal was normalized to Tubulin signal in individual experiments and was set to 1 in wild-type. D) Q-PCR analysis of LTR transposons *burdock*, *blood* and *gypsy* and non-LTR transposon *HeT-A* RNAs in *vas^D1/D1^*; *nos-Gal4*>GFP-Vas^WT^ *and vas^D1/D1^*; *vas-Gal4*>GFP-Vas^WT^ early embryos. Expression level of transposons in wild-type (*w^1118^)* was set to 1 and normalized to *18S* rRNA in individual experiments. Error bars represent the standard deviation among three biological replicates. Student’s *t-*test indicated *p*-values for *burdock* (0.004), *blood* (0.002) and *HeT-A* (0.008) lower than 0.01 (indicated in the chart), while *gypsy* levels were not significantly different (*p* = 0.4). E) Western blot analysis using antibodies against HeT-A/Gag showing protein levels in early embryos produced by wild-type (*w^1118^*), *vas^D1/D1^*; *nos-Gal4*>GFP-Vas^WT^ *and vas^D1/D1^*; *vas-Gal4*>GFP-Vas^WT^ flies. Tubulin was used as a loading control. Table shows quantification of HeT-A/Gag protein levels relative to wild-type. HeT-A/Gag signal was normalized to Tubulin signal in individual experiments and was set to 1 in wild-type.

The correlation between high levels of transposon expression during oogenesis (Figure 1A, *nos-Gal4*-driven) and a high frequency of nuclear damage and DSBs in *vas^D1/D1^*; *nos-Gal4*>GFP-Vas^WT^ embryos (Figure 2A-C) suggested that maternally transmitted transposons cause embryonic lethality. To test this we compared transposon RNA levels in embryos of *vas^D1/D1^*; *nos-Gal4*>GFP-Vas^WT^ and *vas^D1/D1^*; *vas-Gal4*>GFP-Vas^WT^ flies, in which transposon RNAs are up- and down-regulated, respectively (Figure 1A). Levels of maternally transmitted transposon RNA were significantly higher in *vas^D1/D1^*; *nos-Gal4*>GFP-Vas^WT^ embryos (Figure 2D) suggesting that the increased lethality observed in *vas^D1/D1^*; *nos-Gal4*>GFP-Vas^WT^ embryos is due to DNA damage (Figure 2A-C) caused by the high levels of maternally transmitted transposon RNAs (Figure 2D)

One of the up-regulated transposons in *vas* mutants is *HeT-A*, whose RNA and protein expression is strongly de-repressed in piRNA pathway mutant ovaries (Aravin et al., 2001; Lopez-Panades et al., 2015; Vagin et al., 2006; Zhang et al., 2014). Analysis of HeTA/Gag protein expression in 0-1 hour old embryos showed that levels of HeT-A/Gag were much higher in *vas^D1/D1^*; *nos-Gal4*>GFP-Vas^WT^ than in *vas^D1/D1^*; *vas-Gal4*>GFP-Vas^WT^ embryos (Figure 2E). Additionally, we stained embryos with antibodies against HeT-A/Gag protein and observed that in cellularized wild-type embryos, HeT-A localized at distinct perinuclear foci (Figure 3A panel a and Supplementary Figure S3B panel a), as previously described for HeT-A/Gag-HA-FLAG fusion protein (Olovnikov et al., 2016). In *vas^D1/D1^*; *nos-Gal4*>GFP-Vas^WT^ embryos displaying nuclear damage HeT-A protein accumulated in large foci throughout the embryo (Figure 3A panel b and Supplementary Figure S3B panel b), whereas embryos of the same genotype lacking nuclear damage showed a wild-type distribution of the protein (Figure 3A panel c and Supplementary Figure S3B panel c). Finally, HeT-A/Gag displayed wild-type localization in *vas^D1/D1^*; *vas-Gal4>*GFP-Vas^WT^ embryos (Figure 3A panel d and Supplementary Figure S3B panel d). Altogether, these results show that up-regulation of transposon mRNAs and proteins during oogenesis results in their maternal transmission to the progeny, where they cause DSBs, nuclear damage, and arrest of embryogenesis.

**Figure 3:**
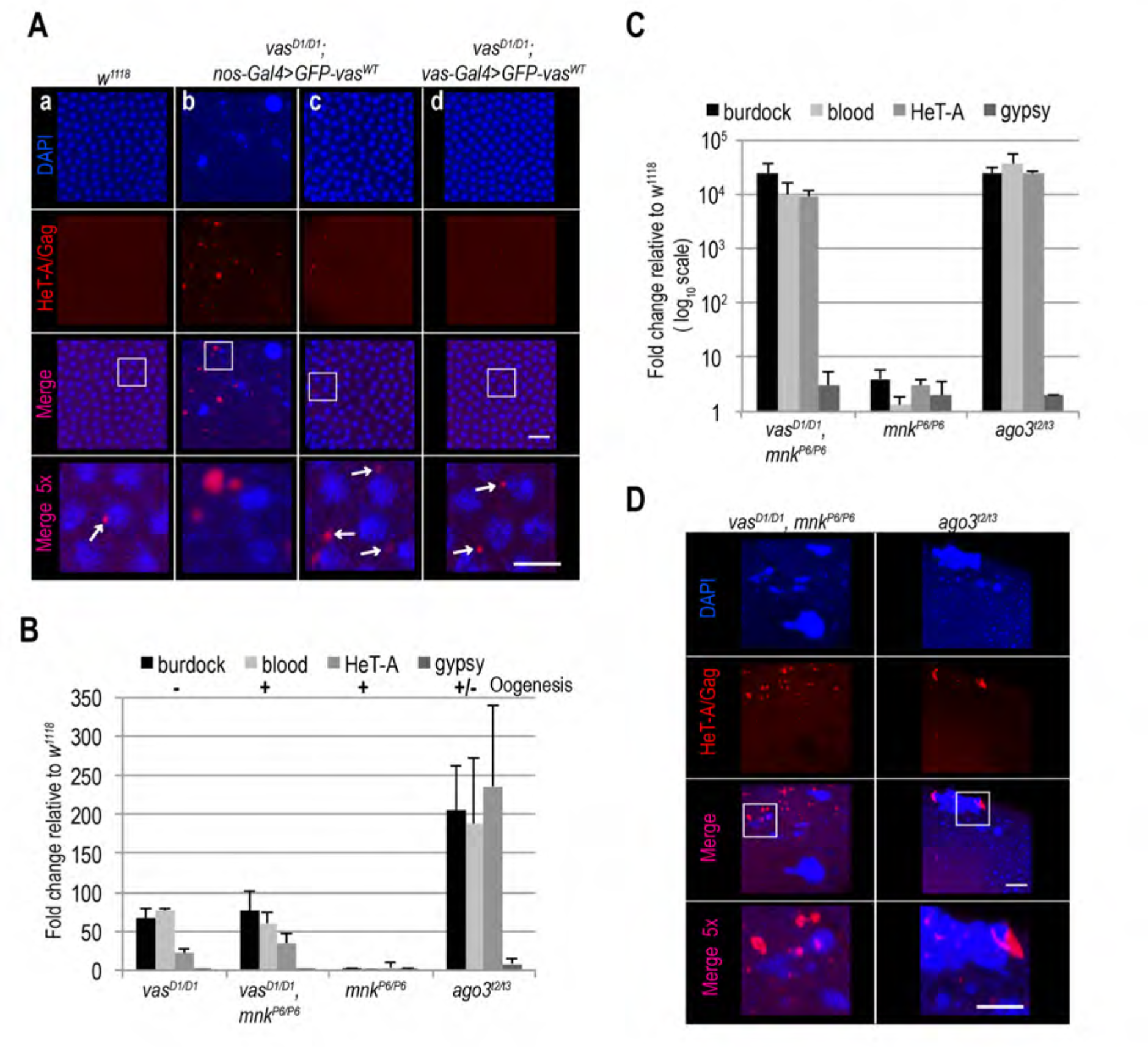
Loss of Chk2 DNA damage signaling does not restore embryogenesis in *vas* mutant flies. A) Immunohistochemical detection of HeT-A/Gag protein in wild-type (*w^1118^*; a*), vas^D1/D1^*; *nos-Gal4*>GFP-Vas^WT^ (b and c) and *vas^D1/D1^*; *vas-Gal4*>GFP-Vas^WT^ (d) stage 5 embryos. Arrows indicate wild-type localization of HeT-A/Gag. Staining of the whole embryos is presented in Supplementary Figure S3A. Scale bars indicate 10 µm and 5 µm (5x magnification). B) Q-PCR analysis of LTR transposons *burdock*, *blood* and *gypsy* and non-LTR transposon *HeT-A* RNAs in ovaries from *vas^D1/D1^* single and *vas^D1/D1^, mnk^P6/P6^* double mutant flies, as well as *mnk^P6/P6^* and *ago^t2/t3^* mutant flies. Expression level of transposons in wild-type (*w^1118^)* was set to 1 and normalized to *rp49* mRNA in individual experiments. Error bars represent the standard deviation among three biological replicates. Oogenesis completion is indicated with + and -. C) Q-PCR analysis of LTR transposons *burdock*, *blood* and *gypsy* and non-LTR transposon *HeT-A* RNAs in early embryos produced by *vas^D1/D1^, mnk^P6/P6^* double mutant, and *mnk^P6/P6^* and *ago^t2/t3^* single mutant flies. Expression level of transposons in wild-type (*w^1118^)* was set to 1 and normalized to *18S* rRNA in individual experiments. Error bars represent the standard deviation among three biological replicates. D) Immunohistochemical detection of HeT-A/Gag protein in stage 5 embryos from *vas^D1/D1^*, *mnk^P6/P6^* double mutant and *ago^t2/t3^* single mutant flies. Staining of the whole embryos is presented in Supplementary Figure S4D. Scale bars indicate 10 µm and 5 µm (5x magnification).

### Chk2 mutation restores oogenesis but not transposon silencing and embryogenesis in *vas* mutants

To test genetically whether DNA damage signaling contributes to the oogenesis arrest of *vas* loss-of-function mutants (Hay et al., 1988; Lasko and Ashburner, 1990; Schüpbach and Wieschaus, 1986 and Figure 1A), we introduced the *mnk^P6^* loss-of-function allele into the *vas^D1^* background. Genetic removal of *mnk* (Supplementary Figure S3B) suppressed the oogenesis arrest of *vas^D1/D1^* mutants and partially rescued their egg-laying (Supplementary Figure S3C and S4A). Importantly, *mnk^P6/P6^* single mutants expressed Vasa at wild-type levels and, as expected, the protein was not detected in *vas^D1/D1^, mnk^P6/P6^* double mutants (Supplementary Figure S2C). Taken together, these findings demonstrate that the oogenesis arrest of loss-of-function *vas* mutants results from activation of the Chk2-mediated DNA damage-signaling checkpoint.

Although removal of *mnk* allowed oogenesis progression, it did not reduce transposon levels in *vas^D1/D1^, mnk^P6/P6^* ovaries and the eggs laid failed to hatch (Figure 3B and Supplementary Figure S4B). Further analysis revealed that *vas^D1/D1^, mnk^P6/P6^* early embryos contained elevated levels of maternally transmitted transposon RNAs (Figure 3C). This was also the case of *ago3* single mutant embryos, which displayed nuclear damage (Mani et al., 2014; Supplementary Figure S4D) similar to that of *vas^D1/D1^*; *nos-Gal4*>GFP-Vas^WT^ embryos (Figure 2A-B). In addition to HeT-A RNA, HeT-A/Gag protein was also up-regulated in *vas^D1/D1^, mnk^P6/P6^* and *ago3* embryos during the syncytial blastoderm stage (Supplementary Figure S4C). At cellularization, the embryos displayed nuclear damage and HeT-A/Gag was present in large foci throughout the embryo (Figure 3D and Supplementary Figure S4D), resembling the nuclear-damaged *vas^D1/D1^*; *nos-Gal4*>GFP-Vas^WT^ embryos (Figure 3A panel b).

We next examined the distribution of *HeT-A* RNAs and occurrence of DNA DSBs by fluorescent *in situ* hybridization (FISH) and antibody staining of γH2Av, respectively. Damaged nuclei in *vas^D1/D1^, mnk^P6/P6^* embryos were γH2Av-positive (Figure 4A and Supplementary Figure S5A), indicating that DNA DSBs cause nuclear defects. HeT-A RNAs localized in large foci in *vas^D1/D1^, mnk^P6/P6^* embryos, and was not detectable in wild-type embryos (Figure 4A and Supplementary Figure 5A). Although we did not detect *HeT-A* transcripts in the damaged nuclei of *vas^D1/D1^, mnk^P6/P6^* embryos, the oocyte nucleus was positive both for *HeT-A* RNA and gH2Av (Figure 4B), indicating the presence of DNA DSBs. Further analysis showed that *HeT-A* RNA and HeT-A/Gag protein co-localize in the oocyte cytoplasm and nucleus (Figure 4C) indicating that transposon insertions into the maternal genome begin already during oogenesis. Additional FISH analyses showed that in wild-type egg-chambers, *HeT-A* and *Burdock* transcripts were only detected at sites of transcription in the nurse cell nuclei, while in *vas^D1/D1^, mnk^P6/P6^* and *^ago3^* egg-chambers transcripts of both transposons accumulated within the oocyte along the anterior margin, as well as around and within the nucleus (Figure 4D and Supplementary Figure S5B). These results show that in *vas^D1/D1^, mnk^P6/P6^* double and in *ago3* single mutant females up-regulated transposons invade the maternal genome and are transmitted to the progeny, causing severe nuclear defects and embryogenesis arrest. We conclude that tight regulation of transposons throughout oogenesis is essential to maintain genome integrity in the oocyte and in early syncytial embryo, hence for normal embryonic development.

**Figure 4:**
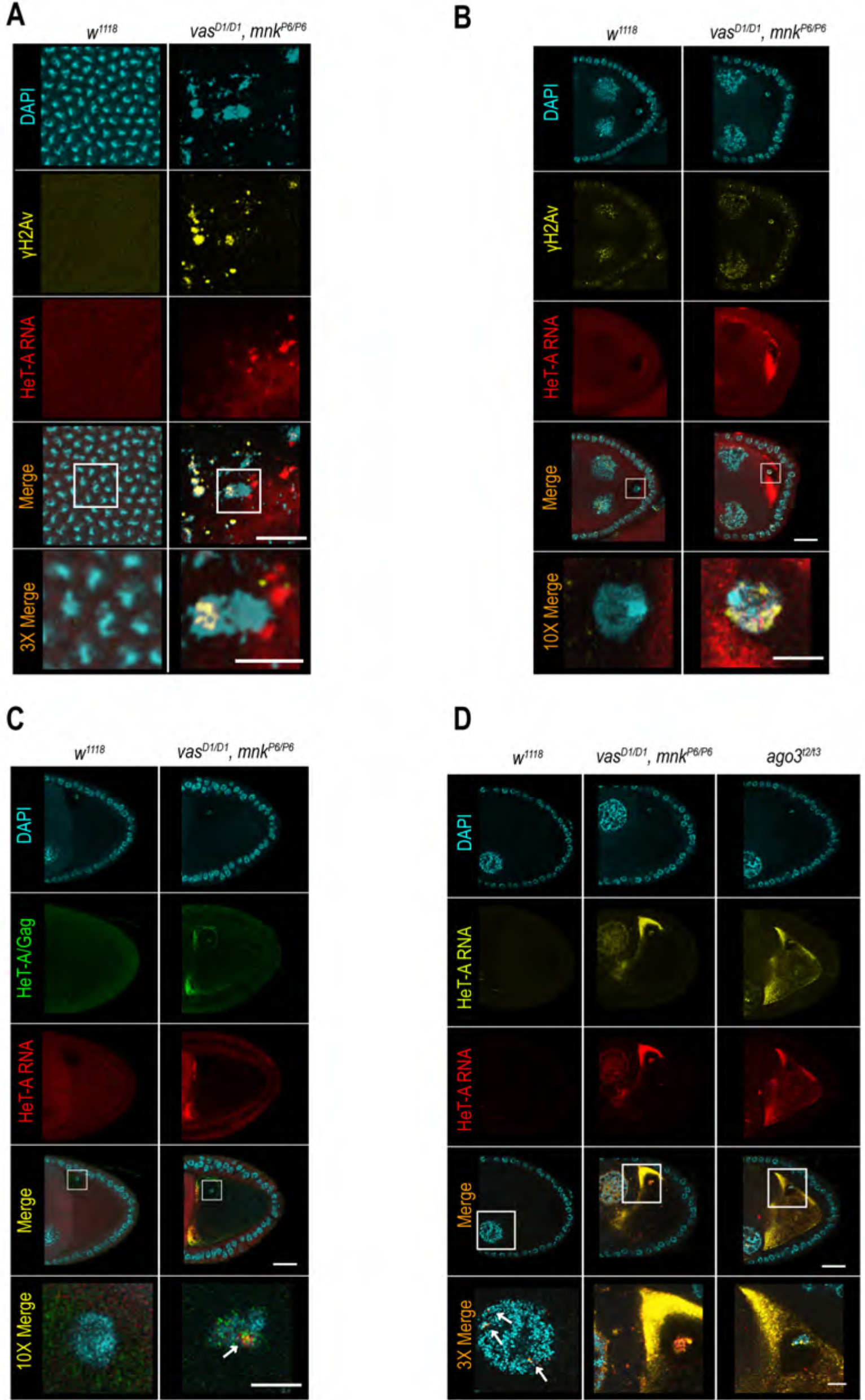
Transposons invade maternal genome and cause DNA DSBs in *vas, mnk* double mutant flies. A and B) *In situ* detection of *HeT-A* mRNA by FISH and immunohistochemical detection of DNA double-strand breaks using antibodies against H2Av pS137 (γH2Av) in wild-type (*w^1118^*) and *vas^D1/D1^, mnk^P6/P6^* double mutant embryos (A) and ovaries (B). Scale bars in (A) indicate 5 µm and 2 µm (3x magnification); scale bars in (B) indicate 20 µm and 5 µm (10x magnification). C) *In situ* detection of *HeT-A* mRNA by FISH and immunohistochemical detection of HeTA/Gag protein in wild-type (*w^1118^*) and *vas^D1/D1^, mnk^P6/P6^* double mutant ovaries. Arrow indicates co-localization of *HeT-A* mRNA and HeT-A/Gag protein signals. Scale bars indicate 20 µm and 5 µm (10x magnification). D) *In situ* detection of *HeT-A* mRNA by FISH in wild-type (*w^1118^*), *vas^D1/D1^, mnk^P6/P6^* double mutant and *ago^t2/t3^* single mutant ovaries. Arrows indicate sites of *HeT-A* mRNA transcription. Scale bars indicate 20 µm and 5 µm (3x magnification).

## Discussion

Our study shows that a transient loss of *vas* expression during early oogenesis leads to up-regulation of transposon levels and compromised viability of progeny embryos. The observed embryonic lethality is due to DNA DSBs and nuclear damage that arise as a consequence of the elevated levels of transposon mRNAs and proteins, which are transmitted from the mother to the progeny. We thus demonstrate that transposon silencing in the nurse cells is essential to prevent maternal transmission of transposons and DNA damage, protecting the progeny from harmful transposon-mediated mutagenic effects.

Our finding that suppression of Chk2-mediated DNA damage signaling in loss-of-function *vas* mutant flies restores oogenesis and egg production demonstrates that Chk2 is epistatic to *vas*. However, hatching is severely impaired, due to the DNA damage sustained by the embryos. The defects displayed by *vas, mnk* double mutant embryos resembled those of PIWI (*piwi*, *aub* and *ago3*) single and *mnk*; PIWI double mutant embryos (Klattenhoff et al., 2007; Mani et al., 2014). Earlier observation that inactivation of DNA damage signaling does not rescue the development of PIWI mutant embryos led to the assumption that PIWI proteins might have an essential role in early somatic development, independent of cell cycle checkpoint signaling (Mani et al., 2014). By tracing transposon protein and RNA levels and localization from the mother to the early embryos we have shown that, independent of Chk2 signaling, de-repressed transposons are responsible for nuclear damage and embryonic lethality. Our study demonstrates that transposon insertions occur in the maternal genome where they cause DNA DSBs that together with transposon RNAs and proteins are passed on to the progeny embryos. Transposon activity and consequent DNA damage in the early syncytial embryo cause aberrant chromosome segregation, resulting in unequal distribution of the genetic material, nuclear damage and ultimately embryonic lethality. Our study shows that early *Drosophila* embryos are defenseless against transposons and will succumb to their mobilization if the first line of protection against selfish genetic elements in the nuage of nurse cell fails.

A recent study showed that in *p53* mutants transposon RNAs are up-regulated accumulate at the posterior pole of the oocyte, without deleterious effects on oogenesis or embryogenesis (Tiwari et al., 2017). It is possible that the absence of pole plasm in *vas* mutants (Lehman and Ephrussi, 1994) results in release of the transposon products and their ectopic accumulation in the oocyte. Localization of transposons to the germ plasm (Tiwari et al., 2017) may restrict their activity to the future germline and protect the embryo soma from transposon activity. Transposon-mediated mutagenesis in the germline would produce genetic variability, a phenomenon thought to play a role in the environmental adaptation and evolution of species. It would therefore be of interest to determine the role of pole plasm in transposon control in the future.

Transposon up-regulation in the *Drosophila* female germline triggers a DNA damage-signaling cascade (Chen et al., 2007, Klattenhoff et al., 2007). In *aub* mutants, before their oogenesis arrest occurs, Chk2-mediated signaling leads to phosphorylation of Vasa, leading to impaired *grk* mRNA translation and embryonic axis specification (Klattenhoff et al., 2007). Considering the genetic interaction of *vas* and *mnk* (Chk2) and the fact that Vasa is phosphorylated in Chk2-dependent manner (Abdu et al., 2002, Klattenhoff et al., 2007) it is tempting to speculate that phosphorylation of Vasa might stimulate piRNA biogenesis, reinforcing transposon silencing and thus minimizing transposon-induced DNA damage (Figure 5). The arrest of embryonic development as a first, and arrest of oogenesis as an ultimate response to DNA damage thus prevents the spreading of detrimental transposon-induced mutations to the next generation.

**Figure 5:**
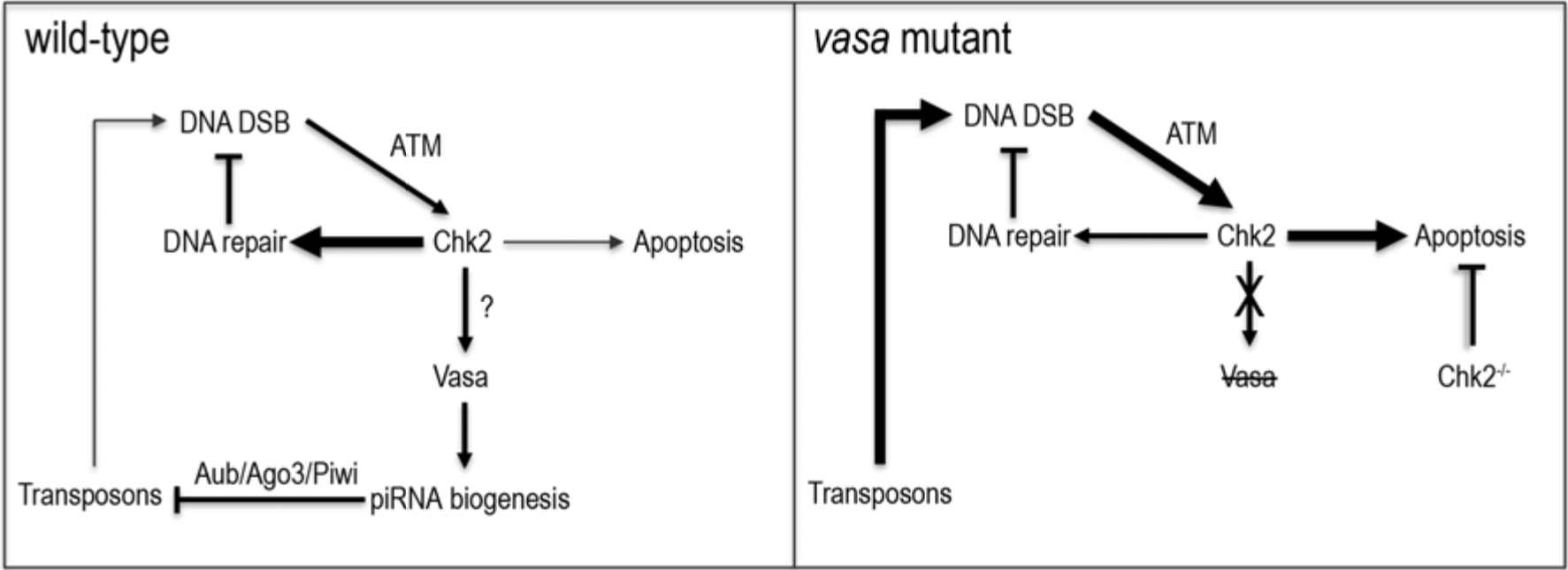
Vasa couples the DNA damage response machinery and the piRNA pathway in *Drosophila* female germline. In wild-type flies, the occurrence of DNA DSBs activates Chk2 kinase that regulates several mechanisms that together antagonize deleterious effects of DNA damage. Chk2 might directly or indirectly target Vasa that in turn affects piRNA biogenesis and transposon control reducing the transposon-induced DSBs. Accordingly, DNA damage induced by high levels of transposons in *vas* mutants triggers DNA damage-induced apoptosis resulting in oogenesis arrest. Oogenesis can be restored by depletion of Chk2 however transposon deregulation persists and causes severe nuclear damage and embryogenesis arrest preventing distribution of transposon-induced, detrimental mutations within the population.

## Experimental Procedures

### Fly stocks and husbandry

The following *Drosophila* stocks were used: *w^1118^*; *b^1^, vas^D1^/CyO* (*vas^3^*, Tearle and Nusslein-Volhard, 1987, Lasko and Ashburner, 1990); *b^1^, vas^Q7^, pr^1^/CyO* (*vas^7^*, Tearle and Nusslein-Volhard, 1987, Lasko and Ashburner, 1990); *vas^D1^/CyO; nos-Gal4-VP16/TM2* (Xiol et al., 2014); *vas-Gal4* (gift of Jean-René Huynh); *GFP-vas^WT^/TM2* (Xiol et al., 2014); *mnk^P6^/CyO* (Brodsky et al., 2004); *bw^1^; st^1^, ago3^t2^/TM6B, Tb^+^* (FBst0028269), *bw^1^; st^1^; ago3^t3^/TM6B, Tb^1^* (FBst0028270). All flies were kept at 25°C on standard *Drosophila* medium.

### Generation of *mnk, vas* double mutant flies

To generate *mnk, vas* double mutants, *+, +, mnk^P6 [P{lacW}]^/CyO* and *b^1^, vas^D1^, +/CyO* flies were crossed. F1 progeny *+, +, mnk^P6 [P{lacW}]^/ b^1^, vas^D1^, +* females were then crossed to males of the balancer stock *CyO/if*. F2 progeny were screened for red eyes (*mnk^P6^* marker *P{lacW})* and 200 individual red-eyed flies were mated to CyO/if balancer flies. F3 generation stocks were established and screened for non-balanced flies of a dark body color (homozygous for *b^1^*, a marker of the original *vas^D1^* chromosome). 3 lines were obtained and tested for presence of the *vas ^D1^* mutation by western blotting (Supplementary Figure S2C) and for presence of the *mnk* mutation by RT-PCR (Supplementary Figure S3B). A scheme of the crosses and recombination is shown in Supplementary Table S1 and sequences of primers used for RT-PCR reaction are shown in Supplementary Table S7.

### Fecundity and hatching assays

Virgin females of all genetic backgrounds tested were mated with *w^1118^* males for 24h at 25°C. The crosses were then transferred to apple-juice agar plates, and eggs collected in 24h intervals over 3-4 days. The number of eggs laid on each plate was counted; the plates were kept at 25°C for 2 days, then the number of hatched larvae counted. Experiments were performed in three independent replicates, and *w^1118^* females were used as a control.

### Ovarian morphology and Vasa localization analysis

Ovaries of 3-7-days old flies were dissected in PBS. Ovarian morphology was evaluated under a Olympus SZX16 stereo microscope. Vasa localization was assessed in ovaries of 3-7-days old flies expressing the GFP-Vasa proteins after fixation in 2% PFA and 0.01% Triton X-100 for 15 min at RT. Fixed ovaries were mounted on glass slides and GFP fluorescence examined under a Zeiss LSM 780 confocal microscope. Vasa localization in wild-type and *vas* mutant ovaries and progeny embryos was analyzed by antibody staining (see below). Nuclei were visualised with NucBlue Fixed Cell Stain (Thermofisher).

### Immunohistochemical staining of ovaries and embryos

Freshly hatched females were mated with wild-type males and kept for 2-3 days on yeast at 25°C prior to dissection. Ovaries were dissected in PBS and immediately fixed by incubation at 92°C for 5 min in preheated fixation buffer (0.4% NaCl, 0.3% Triton X-100 in PBS), followed by extraction in 1% Triton X-100 for 1h at room temperature (RT). Fixed ovaries were incubated with primary antibodies against Vasa (rat; 1:500; Tomancak et al., 1998) or HeT-A/Gag (rabbit 1:100; gift of Elena Casacuberta). The following secondary antibodies were used: Alexa 488 conjugated goat anti-rabbit (1:1000; Invitrogen) and Alexa 647 conjugated donkey anti-rat IgG (1:1000; Jackson ImmunoResearch). Nuclei were stained with NucBlue Fixed Cell Stain (Thermofisher).

For embryo staining, freshly hatched females were mated with wild-type males and fed with yeast for 2-3 days at 25°C prior to egg collection. Embryos (0-1h or 1-3h) were collected and dechorionated in 50% bleach, then fixed by incubation at 92°C for 30 sec in preheated fixation buffer (0.4% NaCl, 0.3% Triton X-100 in PBS), followed by devitellinization by rigorous shaking in a 1:1 mix of heptane and methanol. After washing in 0.1% Tween-20, embryos were either immediately incubated with primary antibodies against Vasa (rat; 1:500; Tomancak et al., 1998) or HeT-A/Gag (rabbit 1:100; gift from Elena Casacuberta), or stored in methanol at −20°C for staining later on. For detection of double-strand breaks, embryos (1-3h) were collected and dechorionated in 50% bleach, fixed for 25 min at RT in the heptane/4% formaldehyde interface and devitellinized by rigorous shaking after adding 1V of methanol. After washing in 0.1% Tween-20, the embryos were either immediately incubated with primary antibodies against H2Av pS137 (γH2Av; rabbit; 1:5000; Rockland) or stored in methanol at 20°C for later staining. The following secondary antibodies were used: Alexa 488 conjugated goat anti-rabbit (1:1000; Invitrogen), Alexa 647 conjugated donkey anti-rat IgG (1:1000; Jackson ImmunoResearch) and Alexa 647 conjugated goat anti-rabbit IgG (1:1000; Invitrogen). Nuclei were stained with NucBlue Fixed Cell Stain (Thermofisher). The samples were observed using a Zeiss LSM 780 or Leica SP8 confocal microscope.

### Fluorescent *in situ* RNA hybridization

All FISH experiments were performed as described in Gaspar et al., 2018. In brief, ovaries were dissected in PBS and immediately fixed in 2% PFA, 0.05 % Triton X-100 in PBS for 20 min at RT. Embryos (1-3h) were collected and dechorionated in 50% bleach, fixed for 25 min at RT in the heptane/2% PFA interface and devitellinized by vigorous shaking after adding 1V of methanol. After washing in PBT (PBS + 0.1% Triton X-100) samples were treated with 2 µg/mL proteinase K in PBT for 5 min and then were subjected to 95°C in PBS + 0.05% SDS for 5 min. Proteinase K treatment was omitted when samples were subsequently to be immunohistochemically stained (see bellow). Samples were prehybridized in 200 µL hybridization buffer (300 mM NaCl, 30 mM sodium citrate pH 7.0, 15 % ethylene carbonate, 1 mM EDTA, 50 µg/mL heparin, 100 µg/mL salmon sperm DNA, 1% Triton X-100) for 10 min at 42°C. Fluorescently labeled oligonucleotides (12.5–25 nM) were pre-warmed in hybridization buffer and added to the samples. Hybridization was allowed to proceed for 2 h at 42°C. Samples were washed 3 times for 10 min at 42°C in pre-warmed buffers (1x hybridization buffer, then 1x hybridization buffer:PBT 1:1 mixture, and then 1x PBT). The final washing step was performed in pre-warmed PBT at RT for 10 min. The samples were mounted in 80% 2,2-thiodiethanol in PBS and analyzed on a Leica SP8 confocal microscope.

For simultaneous FISH and immunohistochemical staining, ovaries and embryos were fixed as described above. Samples were simultaneously incubated with fluorescently labeled oligonucleotides (12.5–25 nM) complementary to *HeT-A* RNA and primary antibodies against γH2Av (rabbit; 1:5000; Rockland) or HeT-A/Gag (rabbit 1:100; gift of Elena Casacuberta) overnight at 28°C in PBT. Samples were washed 2 times for 20 min at 28°C in PBT and subsequently incubated with secondary Alexa 488 conjugated goat anti-rabbit antibodies (1:1000; Invitrogen). The samples were mounted in 80% 2,2-thiodiethanol in PBS and analyzed on a Leica SP8 confocal microscope.

### Labeling of DNA oligonucleotides for fluorescent *in situ* RNA hybridization

Labeling of the oligonucleotides was performed as described in Gaspar et al., 2018. Briefly, non-overlapping arrays of 18-22 nt long DNA oligonucleotides complementary to *HeT-A* or *Burdock* RNA (Supplementary Table S7) were selected using the *smFISHprobe_finder.R* script (Gaspar et al., 2017). An equimolar mixture of oligos for a given RNA was fluorescently labelled with Alexa 565- or Alexa 633-labeled ddUTP using terminal deoxynucleotidyl transferase. After ethanol precipitation and washing with 80% ethanol, fluorescently labeled oligonucleotides were reconstituted with nuclease-free water.

### Protein extraction and western blotting

To generate ovarian lysates, around 20 pairs of ovaries from 3-7-day-old flies were homogenized in protein extraction buffer (25 mM Tris pH 8.0, 27.5 mM NaCl, 20 mM KCl, 25 mM sucrose, 10 mM EDTA, 10 mM EGTA, 1 mM DTT, 10% (v/v) glycerol, 0.5% NP40, 1% Triton X-100, 1x Protease inhibitor cocktail (Roche)). For embryo lysates, 0-1h or 1-3h old embryos were collected from apple-juice agar plates and homogenized in protein extraction buffer. Samples were incubated on ice for 10 min, followed by two centrifugations, each 15 min at 16.000 g. 50-100 μg of total protein extracts were solubilized in SDS sample buffer by boiling at 95°C for 5 minutes, then analysed by SDS polyacrylamide gel electrophoresis (4-12% NuPAGE gel; Invitrogen). Western blotting was performed using antibodies against Vasa (rat; 1:3000; (Tomancak et al., 1998)), HeT-A/Gag (rabbit 1:750; gift from Elena Casacuberta), H2Av pS137 (γH2Av; rabbit; 1:1000; Rockland) and Tub (mouse; 1:10000; Sigma T5168).

Quantification of relative protein expression levels was performed using ImageJ. A frame was placed around the most prominent band on the image and used as a reference to measure the mean gray value of all other protein bands, as well as the background. Next, the inverted value of the pixel density was calculated for all measurements by deducting the measured value from the maximal pixel value. The net value of target proteins and the loading control was calculated by deducting the inverted background from the inverted protein value. The ratio of the net value of the target protein and the corresponding loading control represents the relative expression level of the target protein. Fold-change was calculated as the ratio of the relative expression level of the target protein in the wild-type control over that of a specific sample.

### RNA extraction and quantitative PCR analysis

Total RNA was extracted from ovaries of 3-7-day old flies or 0-1h old embryos using Trizol reagent (Thermofisher). For first-strand cDNA synthesis, RNA was reverse-transcribed using a QuantiTect Reverse Transcription Kit (QIAGEN). Quantitative PCR was performed on a StepOne Real-Time PCR System (Thermofisher) using SYBR Green PCR Master Mix (Thermofisher). Relative RNA levels were calculated by the 2^-ΔΔCT^ method (Livak and Schmittgen, 2001) and normalized to *rp49* mRNA levels for ovaries, and 18S rRNA for embryos. Fold-enrichments were calculated by comparison with the respective RNA levels in *w^1118^* flies. Sequences of primers used for qPCR reaction are shown in Supplementary Table S7.

### Data availability

The authors declare that all data supporting the findings of this study are available within the manuscript and its supplementary files.

## Author contributions

Z.D., R.S.P. and A.E. conceived and designed the experiments. Z.D carried out the experiments and analyzed the data. Z.D. and A.E. wrote the manuscript.

## Acknowledgments

We thank Elena Casacuberta for the gift of antibodies against HeT-A/Gag, and Beat Suter and Jean-René Huynh for fly stocks. We are grateful to Anna Cyrklaff and Alessandra Reversi for their help with experiments. We thank the EMBL Advanced Light Microscopy Core Facility (ALMF) for use of its microscopes. This work was funded by the European Molecular Biology Laboratory (EMBL) and Z.D. by a postdoctoral fellowship from the EMBL Interdisciplinary Postdoc Program (EIPOD) under Marie Curie COFUND actions.

---

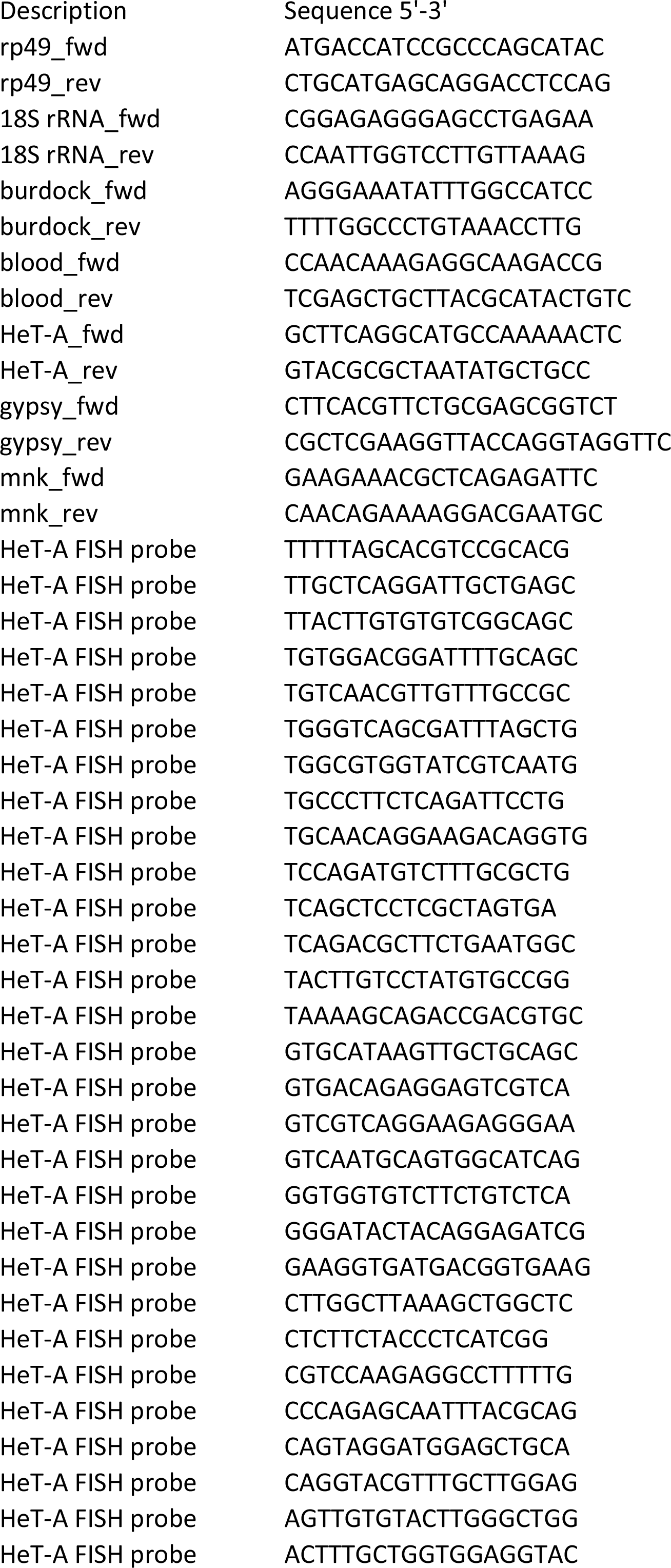
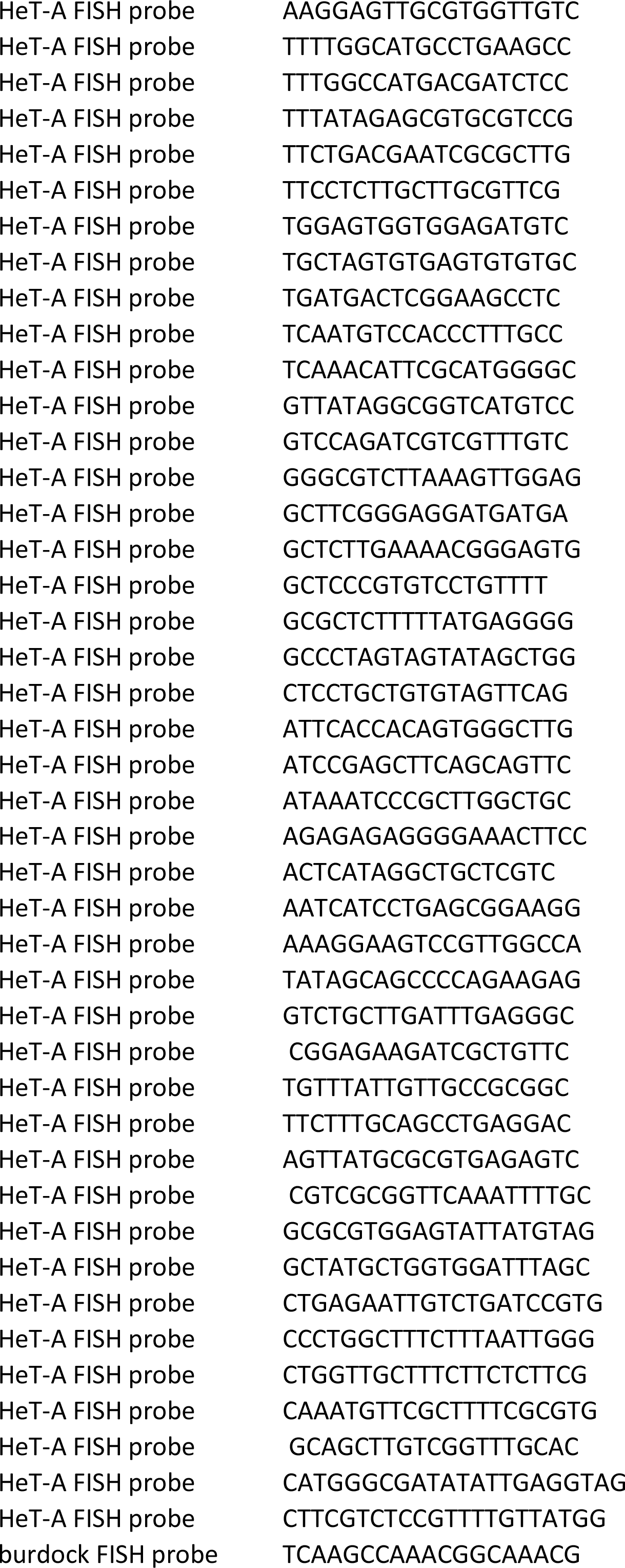
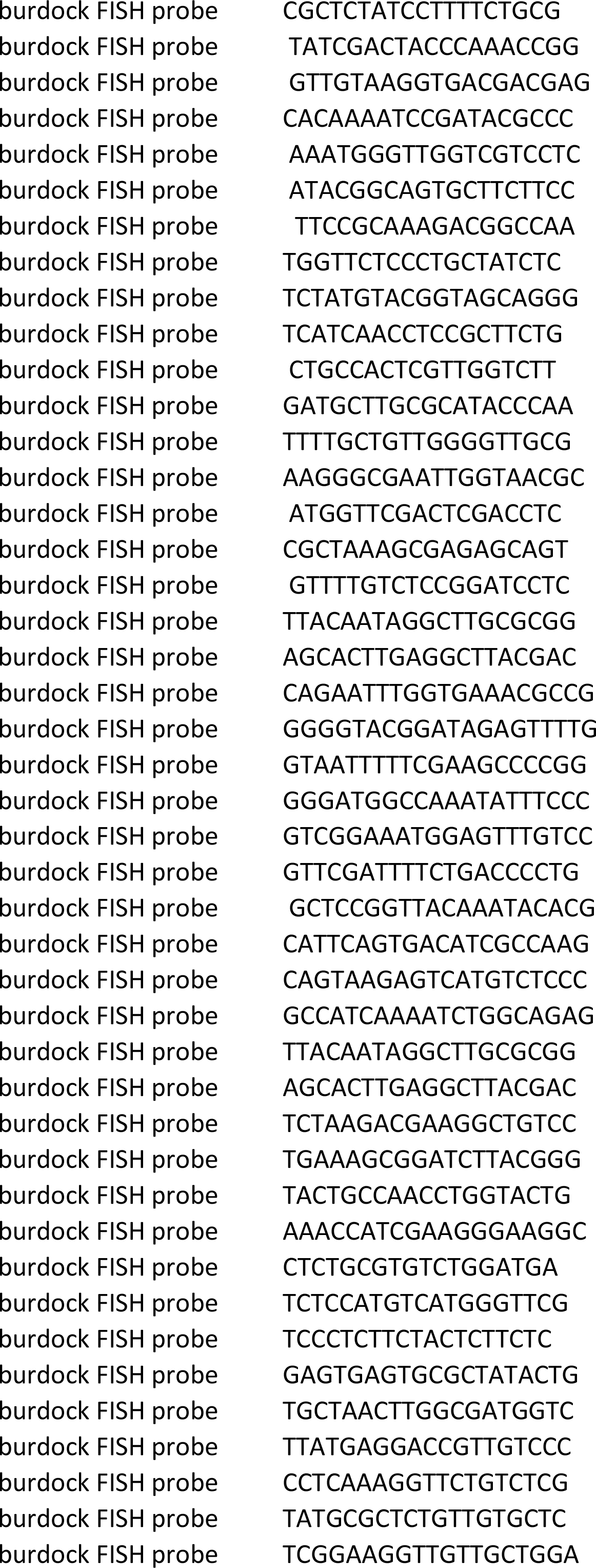
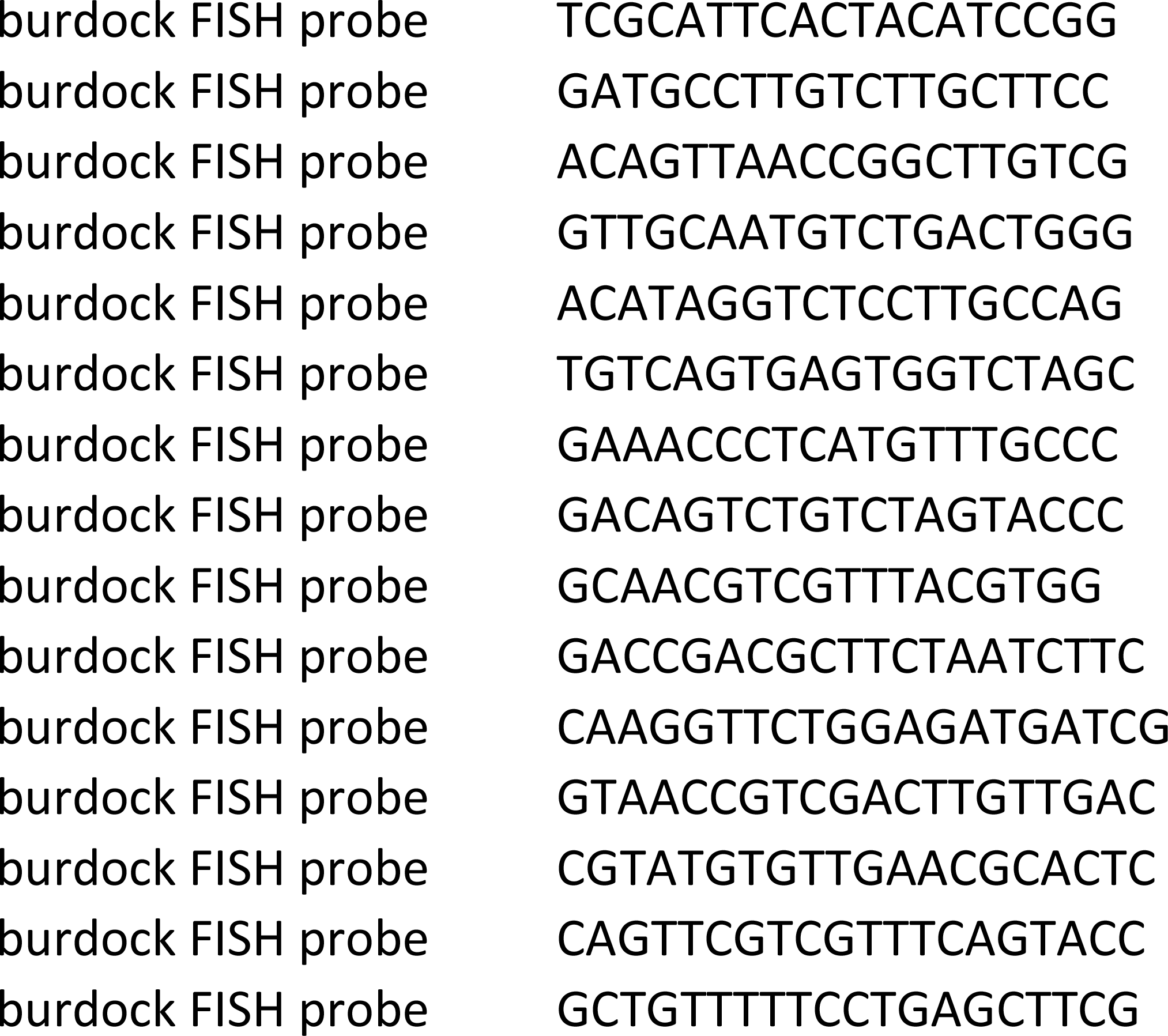

